# Hyaluronidase inhibitor delphinidin inhibits cancer metastasis

**DOI:** 10.1101/2024.04.28.591469

**Authors:** Jeremy McGuire, Taketo Taguchi, Gregory Tombline, Victoria Paige, Michelle Janelsins, Nikesha Gilmore, Andrei Seluanov, Vera Gorbunova

## Abstract

Cancer remains a formidable global health challenge, with metastasis being a key contributor to its lethality. Abundant high molecular mass hyaluronic acid, a major non-protein component of extracellular matrix, protects naked mole rats from cancer and reduces cancer incidence in mice. Hyaluronidase plays a critical role in degrading hyaluronic acid and is frequently overexpressed in metastatic cancer. Here we investigated the potential of targeting hyaluronidases to reduce metastasis. High throughput screen identified delphinidin, a natural plant compound found in fruits and vegetables, as a potent hyaluronidase inhibitor. Delphinidin-mediated inhibition of hyaluronidase activity led to an increase in high molecular weight hyaluronic acid in cell culture and in mouse tissues, and reduced migration and invasion behavior of breast, prostate, and melanoma cancer cells. Moreover, delphinidin treatment suppressed melanoma metastasis in mice. Our study provides a proof of principle that inhibition of hyaluronidase activity suppresses cancer cell migration, invasion and metastasis. Furthermore, we identify a natural compound delphinidin as a potential anticancer therapeutic. Thus, we have identified a path for clinical translation of the cancer resistance mechanism identified in the naked mole rat.

## Introduction

Metastasis, the spread of a primary cancer to a distant site, remains the primary cause of cancer mortality, observed in more than 90% of all cancer deaths ^1-3^. It is imperative that new treatment strategies aimed at preventing metastasis are explored to increase the survival rates for patients with cancer. Metastasis is a complex process that involves multiple steps known as the metastatic cascade, which begins with tumor cells detaching from the primary site, invading local stroma, intravasating into the circulatory or lymphatic system, extravasating at a distant site and growing to form a new tumor ^4,5^. Alteration of the primary tumor microenvironment may affect the initiation of the metastatic cascade and the ability of primary cancer cells to migrate and invade ^6,7^.

Hyaluronic acid (HA), also known as hyaluronan, is a major non-protein component of the extracellular matrix ^7^. HA is a polysaccharide of variable length depending on the number of repeating segments of D-glucuronic acid and N-acetyl-D-glucosamine. The role of HA in cancer progression can be described as a double-edge sword: in some instances, HA can be pro-tumorigenic, but can also be anti-tumorigenic. HA is implicated in cancer progression in multiple cancer types including prostate, bladder, breast and lung ^8-12^. On the other hand, HA is implicated in cancer resistance in the naked mole rat ^13^ as well in tumor suppression of human breast cancer cells ^14^. We recently demonstrated that mice overexpressing hyaluronan synthase 2 from the naked mole-rat show reduced incidence of spontaneous and induced cancers ^15^. An explanation for the dual nature of HA in cancer progression is that high molecular weight hyaluronic acid (HMW-HA) is tumor protective, while low molecular weight hyaluronic acid (LMW HA) is tumor promotive ^16,17^. LMW-HA promotes angiogenesis and inflammation which are all hallmarks of cancer metastasis. ^5,18^. HMW-HA on the other hand is anti-inflammatory ^19^ and inhibits proliferation and migration of cancer cells ^20^. Hence, a balance between HMW-HA and LMW-HA may control tumor progression and metastasis, with HMW-HA being inhibitory for tumor development and metastasis. In humans there is a family of six hyaluronidases (HYALs), HYAL 1- 6 ^21^. HYAL 1 and 2 are the enzymes primarily responsible for degradation of HA in somatic tissues but HYAL 2 is predominately responsible for cleaving HMW-HAs-producing LMW-HAs that are often overexpressed in the tumor microenvironment ^22^. An additional hyaluronidase is transmembrane 2 (TMEM2) is a cell surface protein that possesses potent hyaluronidase activity and may be involved in degrading HA at the cell surface microenvironment ^23^.

We thus sought to identify small molecule inhibitors of hyaluronidases to increase the endogenous levels of HMW-HA that may inhibit cancer progression and metastasis. Since no potent small molecule inhibitors of hyaluronidase were commercially available, we designed an assay for identification of hyaluronidase inhibitors using fluorescence polarization (FP). We conducted a FP-based high-throughput screening assay and identified a natural plant compound, delphinidin, as an inhibitor of hyaluronidase. Delphinidin is the anthocyanidin that gives a dark blue color to many common fruits and vegetables including blueberries, pomegranates, black beans, black grapes, sweet potatoes, pigmented cabbages and some wines such as Cabernet Sauvignon ^24-26^. In our study we show that delphinidin inhibits hyaluronidase degradation of HA both *in vitro* and increases HA levels *in vivo.* When injected to tumor-bearing mice, delphinidin slowed the progression of metastasis. This combined with lack of adverse effects ^27^ makes delphinidin an attractive compound for prevention or treatment of metastatic cancer.

## Results

### High throughput screening for hyaluronidase inhibitors using fluorescence polarization

Increase in HMW-HA achieved by overexpression of naked mole-rat HAS2 gene reduces cancer incidence and extends lifespan in mice ^15^. HMW-HA may confer antimetastatic effects on both lymphatic and vascular metastasis because it enhances lymphatic vessel integrity ^28^. HMW-HA may also inhibit vascular metastasis when primary aortic smooth muscle vascular cells were treated with HA, LMW-HA promoted proliferation while HMW-HA stalled proliferation in the G1 phase ^29^.” In addition, smaller HA oligomers that are generated during HA degradation appear to stimulate tumor invasion and metastasis and we propose that inhibition of hyaluronidases by delphinidin may also block this effect ^30-32^. To translate these benefits to human patients we set out to identify small molecule inhibitors of hyaluronidases which we hypothesized would slow the cleavage of endogenously produced HA and increase HMW-HA levels in tissues. As HMW-HA and LMW-HA differ only in length, we could not use conventional methods such as fluorescent intensity to detect the changes that hyaluronidases make. We applied the concept of FP measurement to detect the difference between the sizes of HA. Conventionally, FP is used to detect the binding of a fluorescent ligand to an enzyme of interest: When a fluorescent ligand is free-floating, it tumbles rapidly. When a polarized light is used to excite the fluorophore, the polarity of emission light would be different from that of excitation light because of the rapid molecular tumbling. On the other hand, when the fluorescent ligand binds to the enzyme of interest, large size of the enzyme prevents it from tumbling rapidly, and after excitation with a polarized light, the polarity of emission light stays close to that of the excitation light. FP measures this similarity between the polarities (the more similar the polarities are, the larger the FP values would be). Fluorescently labeled HMW-HA does not tumble rapidly because of its large size, thus giving large FP values. In contrast, fluorescein labeled LMW-HA tumbles more rapidly and gives smaller FP values. To fluorescently label HA, fluorescein was chosen as a fluorophore because of its appropriate fluorescent lifetime, and the Ugi reaction was used to link fluorescein to HA to achieve zero-length crosslinking and restrict molecular rotation of the fluorophore. By using fluorescent HA and FP, the hyaluronidase inhibitor assay produced reliable measurements with Z’-factor of 0.69 (**Figure 1A)**. Using this assay, about 3000 compounds from the National Cancer Institute (Approved Oncology Drugs Set, Diversity Set, Natural Products Set and Mechanistic Set) were screened (**Figure 1B)**, and Prodelphiniline (**Figure 1C)** was selected as a hit compound. After studying the pharmacophores and structure-activity relationship, a much smaller compound, Delphinidin **(Figure 1D)**, was identified as a small molecule hyaluronidase inhibitor.

**Figure 1.**
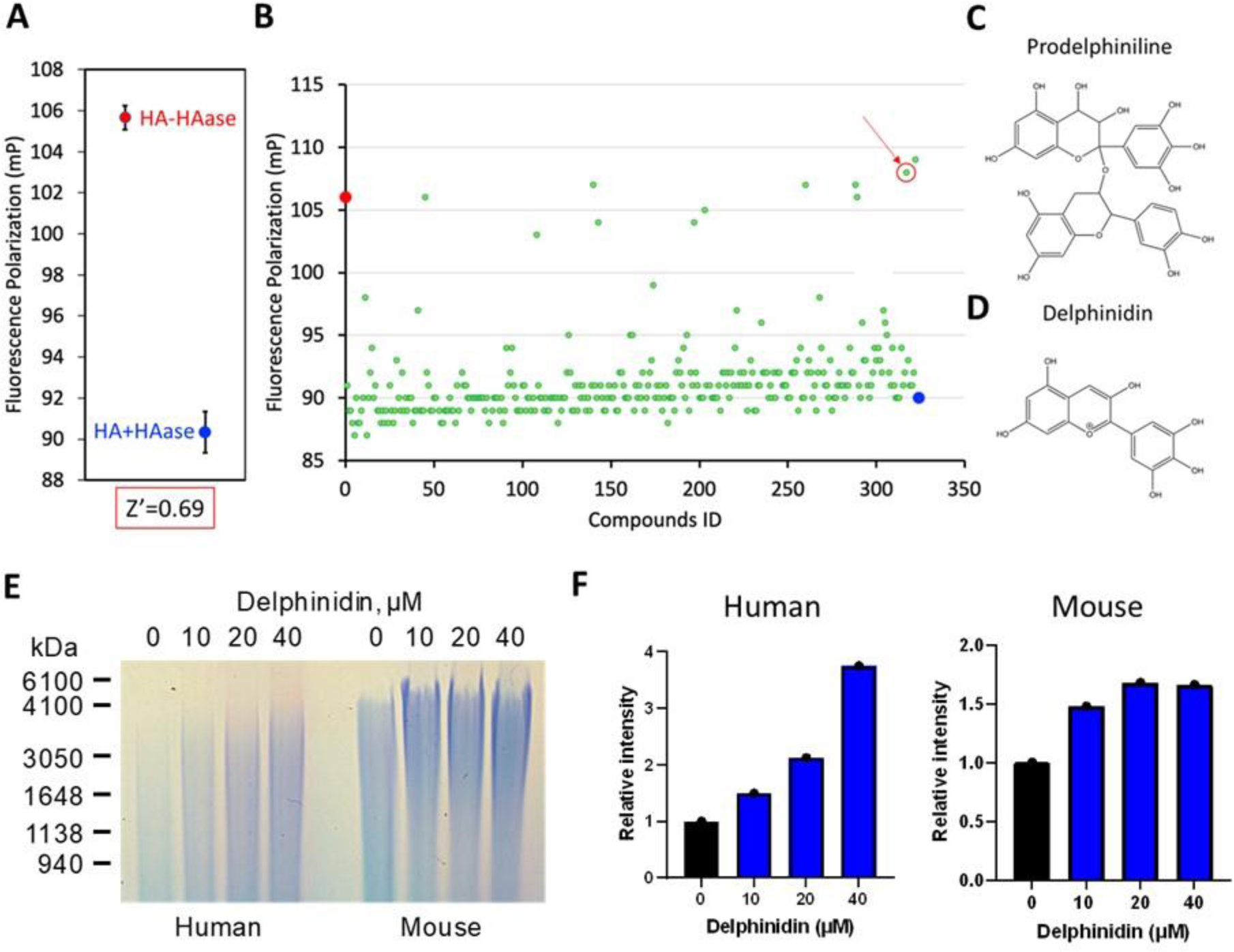
High throughput screening (HTS) for HAase inhibitor using fluorescence polarization & increases in HMW HA by delphinidin. (A) FP shows fluorescent HMW-HA at a higher FP value than LMW-HA (B) Representative data from HTS in a 384 well plate, red arrow indicates prodelphiniline. (C, D) Structures of Prodelphiniline and Delphinidin. (E) Equal numbers of human or mouse fibroblasts were incubated with increasing concentrations of Delphinidin for 24 h. HA was separated on agarose gel (F) Quantification of HA in mouse and human samples.

### Delphinidin inhibits hyaluronidase activity and increases the levels and size of HMW-HA

To test whether delphinidin increases the amount of HMW-HA in cultured cells, mouse and human fibroblasts were treated with delphinidin (0, 10 µM, 20 µM and 40 µM) for 24 hours before hyaluronic acid (HA) was extracted from cell culture media and run on an agarose gel. The size and amount of HA increased in both mouse and human fibroblasts **(Figure 1E)**. The relative concentration of HA was quantitated showing a dose dependent increase in total HA concentration in the delphinidin treated fibroblasts with an increase in HMW HA between 4000 and 600kD observed in the gel image (**Figure 1F**). We extracted HMW-HA from B16-F10 cell culture treated with and without delphinidin. Delphinidin was able to increase the size and amount of HA extracted after 48 hours of culture and when we repeated the experiment with a diluted sample of HAase delphinidin inhibited the degradation of HA **(Supplemental Figure 1).**

To test the efficacy of delphinidin in increasing HA levels *in vivo* we treated 3-month-old wild type C57BL/6 mice with vehicle (n=3) (10% DMSO/90% PBS) or with 50 mg/kg (n=3) or 100 mg/kg of delphinidin (n=3). Intraperitoneal (IP) injections of 100 μl of vehicle or delphinidin were administered three times a week for three weeks before the mice were sacrificed and skin, muscle, kidney, heart, lung, duodenum, jejunum and colon were collected. Previous studies have shown that IP injections of delphinidin provides reliable systemic distribution. There were no visible adverse effects in the mice, and they maintained weight comparable to the vehicle controls. Slides with paraffin embedded tissues from each of the samples collected from the mice were stained using hyaluronic acid binding protein (HABP) and a fluorescent streptavidin antibody to detect HA in the tissues. For each sample one section was treated overnight with hyaluronidase to degrade any HA before staining **(Supplemental Figure 2)**. Using fluorescent microscopy six representative images of each sample for all animals were captured **(Figure 2A)**. Imaris software version 9.8 Oxford Instruments (https://imaris.oxinst.com/) was used for the quantification of maximum fluorescent intensity, and any background fluorescence from the hyaluronidase control samples was subtracted. In skin, muscle, heart, kidney and intestines (duodenum, jejunum and colon) fluorescent intensity was significantly increased in animals treated with delphinidin compared to the vehicle control **(Figure 2B)**. Indicating that delphinidin inhibits hyaluronidases and increases HA levels *in vivo*. While we were not able to test the size of the HA in the tissues, the results from our *in vitro* experiments suggest that HA length is also increased.

**Figure 2.**
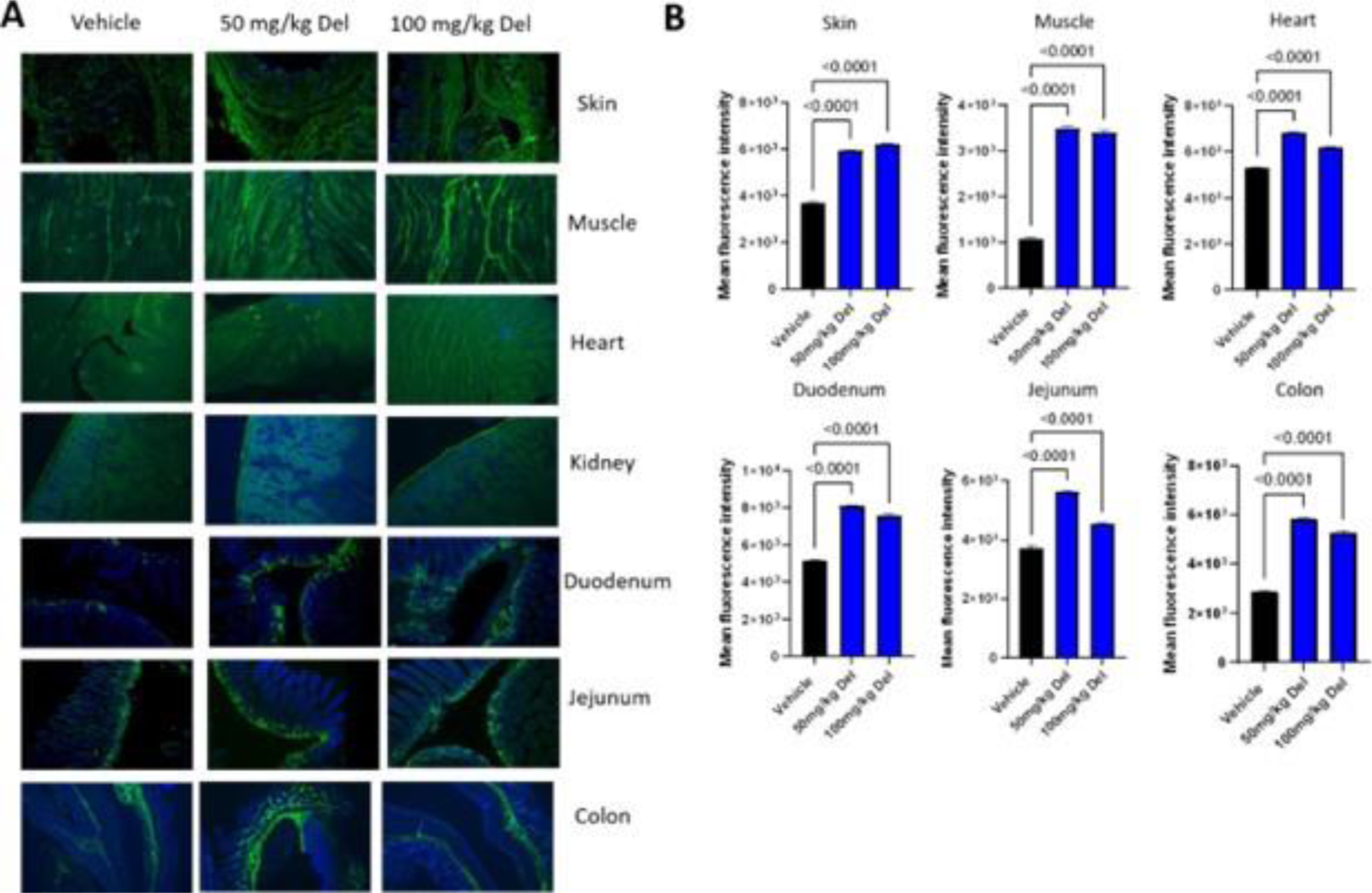
In vivo administration of delphinidin increased HA accumulation in mouse tissues: Mice were injected three times a week for three weeks with either vehicle, 50mg/kg delphinidin or 100mg/kg of delphinidin. Tissues were collected and embedded in paraffin before slide mounting and HABP staining. (A) Representative images (20X) showing HABP staining in mouse tissues. (B) Quantification of HABP staining in mouse tissue mean fluorescence intensity of 6 representative images (three biological replicates with two technical replicates from each tissue) using Imaris cell imaging software. Statistical analyses were generated from one-way ANOVA with multiple comparisons at 95% CI error bars are represented as mean with SEM.

### Delphinidin inhibits cancer cell proliferation

We next tested the ability of delphinidin to inhibit proliferation of normal fibroblasts compared to the three cancer cell lines B16-F10, EO771 and RM1. The rationale for choosing the cell lines used in this study is that they are all metastatic cell lines derived from C57/Bl6 mice and can be used in syngeneic mouse models of metastasis. Mouse breast, prostate and melanoma cancer cells along with wild type mouse fibroblasts were treated with delphinidin or vehicle. Proliferation was quantified using Promega Cell Titer Glo assay 48 hours post treatment. Growth of all three cancer cell lines was inhibited by at least 50% compared to the control when treated with 15 µM delphinidin, whereas 30 µM delphinidin inhibited growth by at least 90%. Skin fibroblasts from wild type mice were not significantly inhibited by 15 µM or 30 µM delphinidin treatment **(Figure 3)**. Delphinidin specifically inhibited cancer cell growth while having little impact on the growth of the normal mouse cells. To test whether the effect of delphinidin is dependent on HA accumulation, the experiment was repeated with the addition of exogenous hyaluronidase (HAase). Cancer cells were treated with either delphinidin alone or delphinidin and exogenous HAase. When HAase was added in combination with delphinidin proliferation was partially restored to all three cancer cell lines **(Figure 3).** These results taken together indicate that delphinidin is inhibiting cancer cell proliferation through the inhibition of HAase.

**Figure 3:**
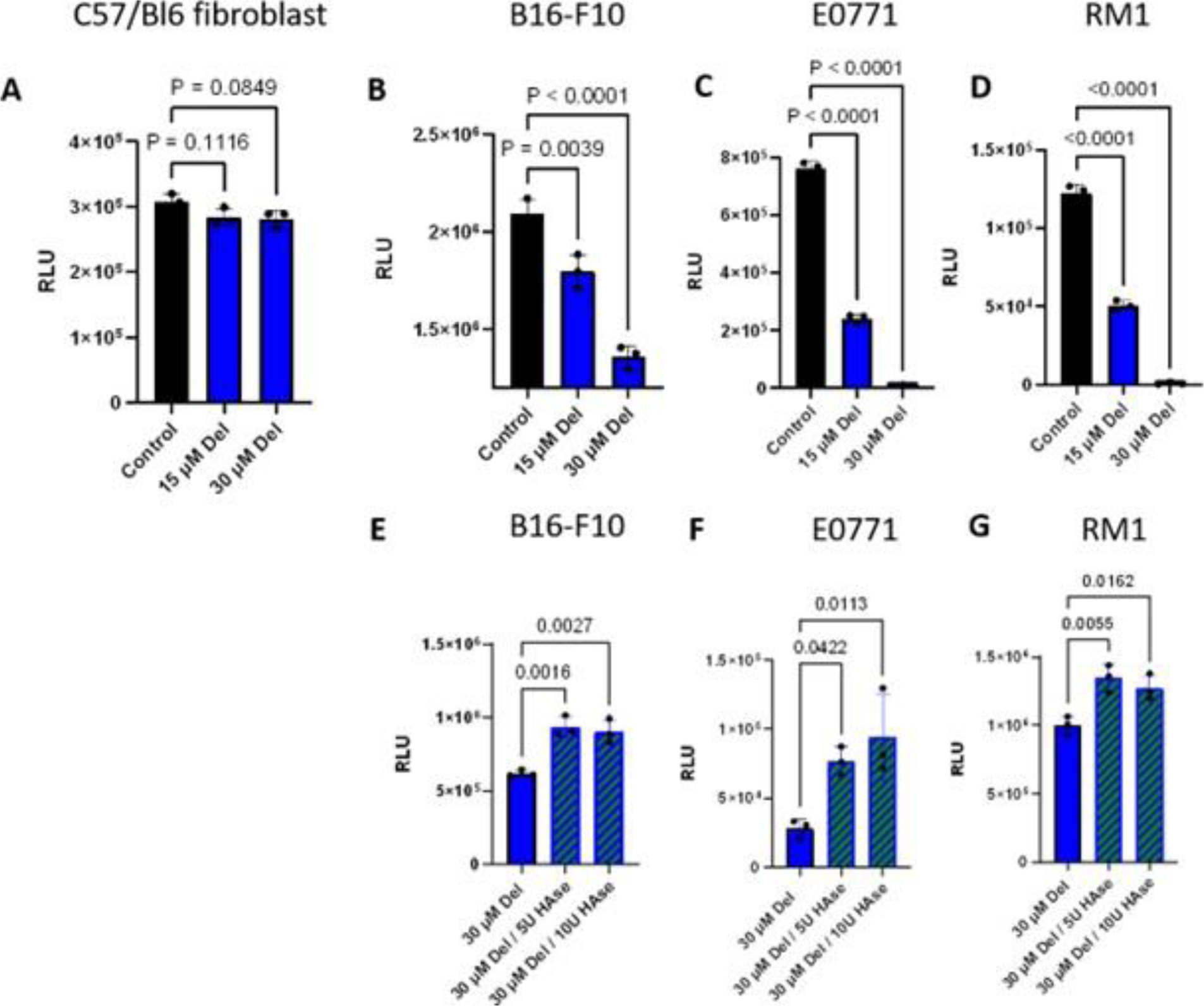
Delphinidin inhibits proliferation of cancer cells. (A) Delphinidin had no significant effect on the growth of wildtype C57/Bl6 fibroblasts after 48 hours. Proliferation was assayed using Cell Titer Glo 2.0 (B-D) Delphinidin in a dose dependent manner significantly inhibited the growth of B16-F10, E0771 and RM1 after 48 hours. (E-G) When the cancer cells were treated with HAase in combination with delphinidin a reduction in growth inhibition was observed at 48 hours. All data was collected as technical triplicates and statistical analyses were generated from one-way ANOVA with multiple comparisons at 95% CI error bars are represented as mean with SEM.

### Delphinidin inhibits migration and invasion of cancer cells

We next assessed if delphinidin has the ability to inhibit both migration and invasion of cancer cell line *in vitro*. Cell migration was first assessed by wound healing assays. Silicone inserts were used to create a gap between the cancer cells so that when the inserts were removed the cells could migrate to fill in the gap. When the cells became confluent the inserts were removed and the cancer cells were treated either with 15 µM delphinidin or control. The cells were then observed hourly over the next 8-20 hours to determine how long it took for the control cells to close approximately 80% of the gap. All three cancer cell lines were significantly inhibited in closing the gap compared to the control cells **(Supplemental Figure 3).**

Cell migration was assessed using a Boyden chamber assay. Cells were seeded in the top chamber, separated from the bottom chamber by a porous membrane. The bottom chambers contained 1% fetal bovine serum (FBS) as a chemoattractant. The top chambers were seeded with cancer cells with or without delphinidin. The addition of 15 µM delphinidin inhibited the migration of all three cancer cell types **(Figure 4)**. Invasion was assessed using a similar Boyden chamber assay the upper chamber is coated with a basement membrane that mimics the *in vivo* microenvironment that cancer cells encounter during the process of metastasis. Cancer cell lines were seeded for 24 hours in triplicate with and without delphinidin. Invasion by all three cancer cell types was significantly inhibited by the addition of 15 µM delphinidin **(Figure 5)**. To test if delphinidin inhibits invasion by its inhibitory effect on HAase, we added delphinidin and HAase to the upper chamber with the cancer cells. The addition of HAase abrogated the inhibitory effect of delphinidin on invasion after 24 hours in all cell lines. **(Figure 5)**. These results suggest that delphinidin inhibits invasion of cancer cells in part by inhibiting HAase.

**Figure 4:**
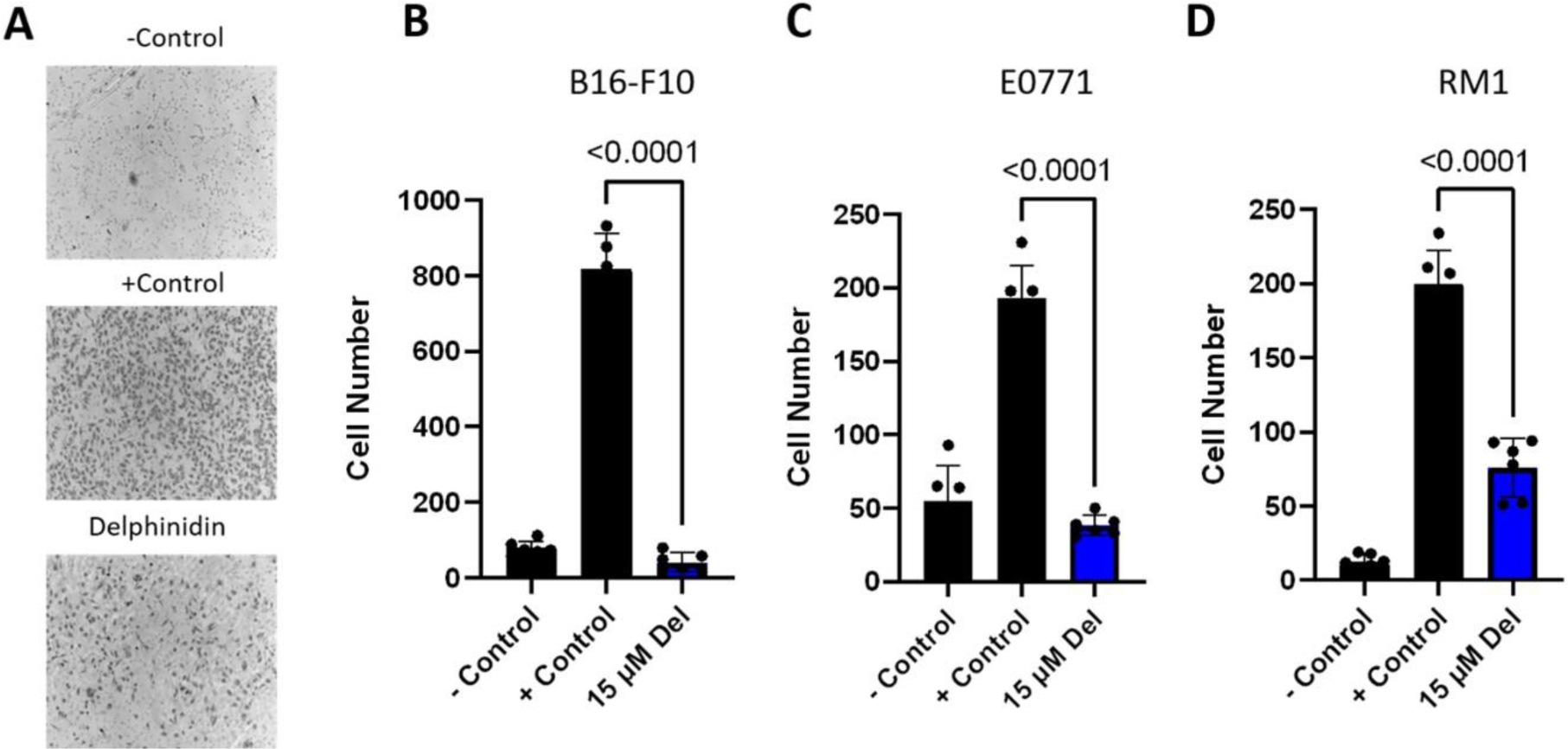
Delphinidin inhibits migration of cancer cells. (A) Representative images of membranes from Boyden chamber assay. Negative control was serum free media on both sides of the membrane and positive control was 1% FBS on the bottom well to promote chemotaxis delphinidin was added to top wells with cells and bottom wells contained 1% FBS (B-D) All three cancer cell lines had significant inhibition of migration when incubated with 15 µM delphinidin. (B) B16-F10 (C) E0771 (D) RM1. All data were collected as technical triplicates and statistical analyses were generated from one-way ANOVA with multiple comparisons at 95% CI error bars are represented as mean with SEM.

**Figure 5:**
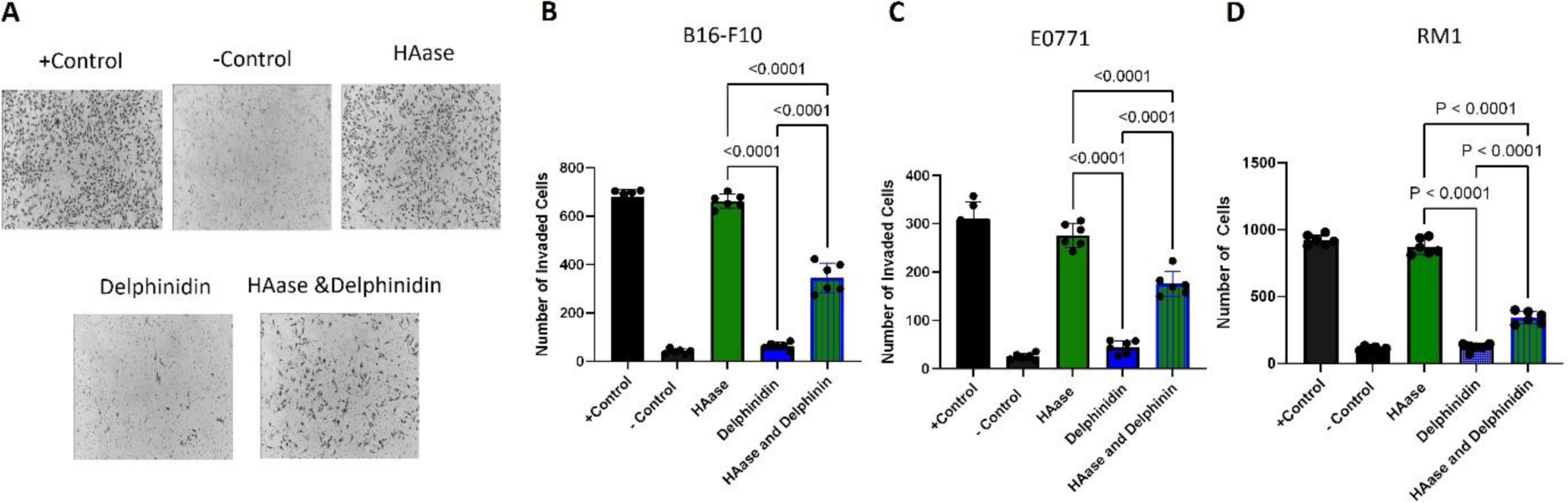
Delphinidin inhibits invasion of cancer cell and HAase abrogates the effect of delphinidin. (A) Representative images of the membranes from the Boyden chamber invasion assay. The inside of the top membrane was coated with basement membrane to mimic metastatic invasion. Negative control was serum free media in top and bottom wells and positive control was 1% FBS on the bottom well to promote chemotaxis delphinidin and HAase was added to top wells with cells in serum free media and bottom wells contained 1% FBS. (B-D) 15 µM Delphinidin inhibited the invasion of all three cancer cell lines compared with 10 U/ml of HAase when cells were incubated with delphinidin in combination with 100 U/ml HAase the amount of inhibition was reduced. (B) B16-F10 (C) E0771 (D) RM1. All data was collected as technical triplicates and statistical analyses were generated from one-way ANOVA with multiple comparisons at 95% CI error bars are represented as mean with SEM.

### Delphinidin inhibits spontaneous melanoma metastasis *in vivo*

To test whether delphinidin inhibits cancer cell metastasis in vivo, 8-month-old C57/BL6 mice were pretreated with either delphinidin 50 mg/kg or vehicle, 10% DMSO/90% PBS for one week three times a week. Mouse melanoma B16-F10 luciferase expressing cells (5X10^5^) were inoculated on the ears of the mice between the skin and the cartilage. This spontaneous metastasis model using ear inoculation of B16-F10 better mimics metastasis than the more commonly used tail vein metastasis model ^28,29^. A total of 12 of the delphinidin treated mice and 5 vehicle treated mice received successful injection as determined by observation of bioluminescent signal on the ear when imaging with the IVIS imager the day following the injections. In vivo imaging was performed weekly to monitor primary tumor growth and detection of metastases and delphinidin treatment was continued at three times a week for the duration of the study **(Figure 6A).** Primary tumor growth was delayed in the delphinidin treated cohort, but the difference did not reach statistical significance **(Figure 6B, C).** Detection of metastases to the sentinel lymph node was observed by week 3 in 80% of the vehicle treated mice and 25% of the delphinidin treated mice **(Figure 6B, D)**. These results indicate that delphinidin strongly inhibits metastatic growth. There was no evidence of any adverse events or side effects associated with systemic delphinidin administration compared to vehicle. There was no significant change in weight of the mice over the course of the experiment **(Supplemental Figure 4).**

**Figure 6:**
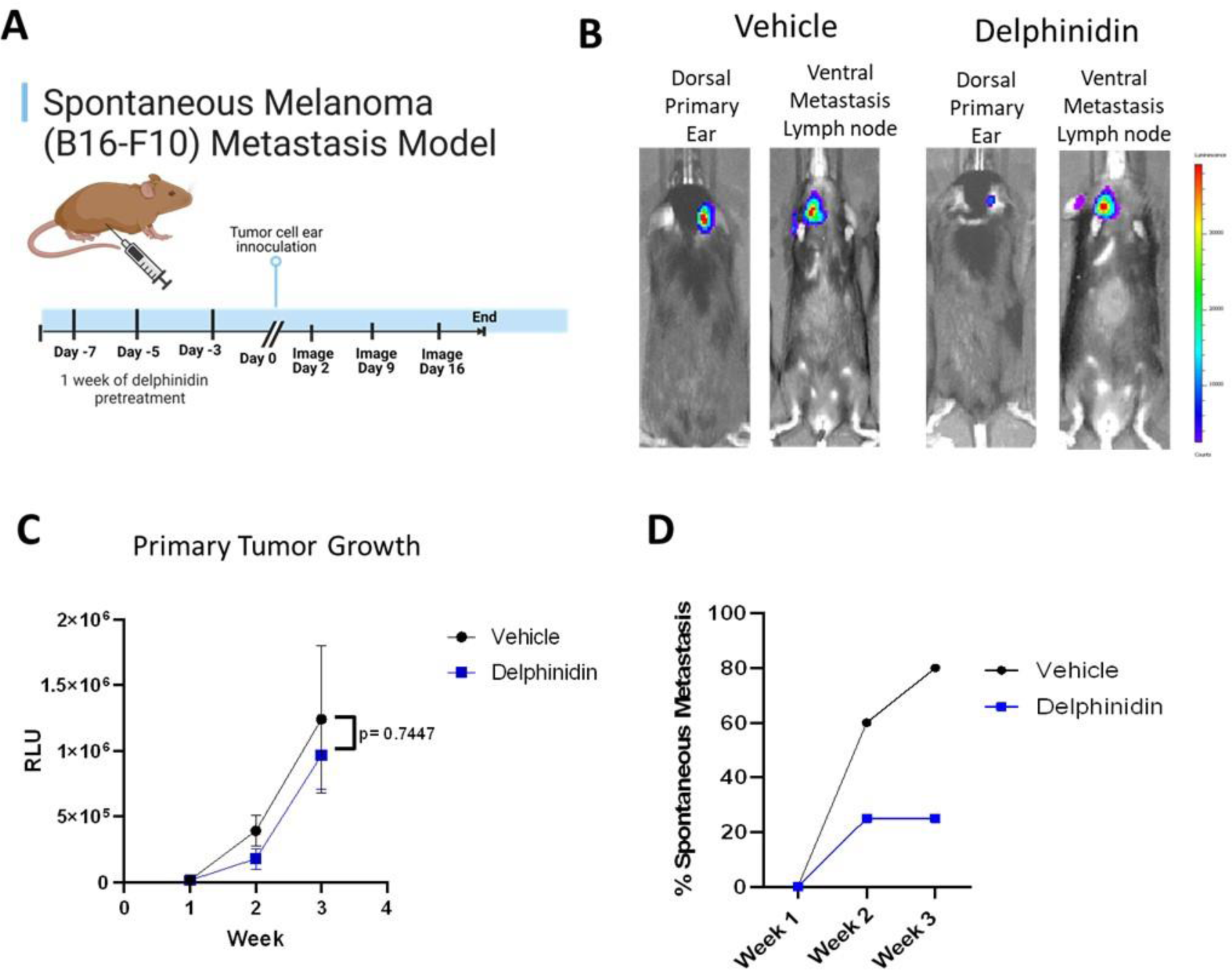
B16-F10 Spontaneous Metastasis model: (A) Model design adapted from “Mouse Experimental Timeline”, by BioRender.com (2023). 50 mg/kg of delphinidin administered by IP 3X a week starting one week before tumor cell (B16-F10) inoculation on the ear and continuing until tumor sizes reached endpoint (2 mm^2^). (B) Representative IVIS images of bioluminescence detection of primary tumor and metastases. (C) Primary tumor growth was not significant between delphinidin treated and vehicle mice. Statistical analysis was generated by unpaired t-test error bars are represented as mean with SEM. (D) At week three (Day 16) 80% (n=4/5) of vehicle treated mice had detectable metastases while only 25% (n=3/12) of delphinidin treated mice had detectable metastases.

## Discussion

While new cancer treatment options have increased survival rates over the past five decades, most cancer deaths are a result of metastatic disease. The metastatic cascade is a complex multistep process that begins with cancer cells invading local stroma and entering the lymphatic or circulatory system. If the initial steps of the metastatic cascade involving cancer cell migration and invasion can be halted, the lives of cancer patients can be extended.

Transgenic overexpression of naked mole-rat HAS2 in mice reduced cancer incidence providing a proof of principle that HMW-HA inhibits cancer *in vivo*. However, the increase in HMW-HA levels achieved in nmrHAS2 expressing mice was modest due to very active hyaluronidase activity in mouse tissues ^15^. As no potent small molecule inhibitors of HAases were available, we performed a chemical screen. One of the challenges in setting up the high-throughput screening assay was that the reactant (HMW-HA) and the product (LMW-HA) are both the same chemical compound: the difference is just its *lengths*. In other words, no chemicals were formed or consumed by the hyaluronidases reaction, and the same amount of "HA" continue to exist before and after the hyaluronidase reaction. We thus could not use conventional methods such as fluorescent intensity to detect the changes that hyaluronidases make. We applied the concept of FP measurement to detect the difference between the sizes of HA, if HA is fluorescently labeled: When HA is HMW-HA, it does not tumble rapidly because of its large size, thus it should give large FP values, whereas LMW-HA should tumble more rapidly and give smaller FP values. Indeed, this FP-based assay allowed us to identify several potential hyaluronidase inhibitors, and the subsequent studies in medicinal chemistry yielded Delphinidin as a small molecule inhibitor for hyaluronidases.

We have shown that delphinidin administered systemically to mice increases hyaluronic acid levels in multiple tissues and may increase levels of tumor protective HMW-HA while preventing the production of tumor supportive LMW-HA in the tumor microenvironment. Delphinidin inhibited proliferation, migration and invasion in melanoma, breast and prostate cancer cells but when delphinidin was administered in combination with exogenous hyaluronidase both proliferation and invasion of cancer cell lines were partially restored. While delphinidin treatment did not significantly impact primary tumor growth of B16-F10 melanoma it did inhibit the percentage of mice that developed spontaneous lymph node metastasis. The ability of delphinidin inhibiting metastasis *in vivo* needs to be confirmed pre-clinically using both spontaneous and experimental metastasis models.

HMW-HA protects the naked mole-rat from cancer and naked mole-rat hyaluronan synthase 2 reduces cancer incidence in transgenic mice ^13,33^. However, elevated levels of HA are found in many cancer types including, lung, breast, prostate and ovary ^34-36^. These elevated HA levels are implicated in causing metastasis and poor prognosis for patients with cancer. This can be explained by concurrent increased hyaluronidase expression in the tumor microenvironment that converts HMW-HA to LMW-HA, giving rise to conflicting data on whether HA is tumor promotive or tumor protective. LMW-HA promotes tumor progression and metastasis while HMW-HA can protect against tumor progression and metastasis ^20^. Elevated levels of hyaluronidase present in the serum of metastatic breast cancer patients compared to patients without metastatic disease may play a role in disease progression ^36,37^ .HA can suppress prostate tumor growth when hyaluronidase is ablated but when HA and hyaluronidase are expressed simultaneously, the result is increased prostate cancer progression and metastasis ^8,37,38^. Thus, the increased amounts of HA that has been associated with tumor progression and metastasis may be due to concurrent elevated levels of the enzyme hyaluronidase degrading HMW-HA to LWM-HA. It should be noted that delphinidin was able to induce apoptosis of the human prostate cancer cell line PC3 ^39^. We tested the ability of delphinidin to induce apoptosis of the mouse cell lines used in our experiments and we did not observe apoptosis using annexin 5 staining in any of the cell lines **(Supplemental Figure 5).** This could be due to different mechanisms of action of delphinidin in human and mouse prostate cancer cells or it could be the concentration of delphinidin (120 and 180 µM) used to induce activation of NF kappa B and apoptosis in the PC3 cell line was much higher than the 30 µM concentration we used in our experiment.

Interestingly, it was previously shown that tea leaf extract that contains delphinidin can inhibit hyaluronidase ^38,40^. As a natural compound delphinidin was reported to have several biological activities many of which may have antitumorigenic potential. Delphinidin was reported to have anti-inflammatory and antioxidant properties ^40-42^. Delphinidin also suppresses VEGF and VEGFR-2, showing anti-angiogenic potential ^42-44^. Delphinidin inhibited metastatic activity of human colorectal cancer cells by inhibiting integrin/FAK cascade ^39,44^, which is consistent with changes in the interaction with extracellular matrix and HA. Delphinidin induces programed cell death, apoptosis, and cell cycle arrest of human prostate cancer cells *in* vitro and *in vivo* ^39,45^. Delphinidin is also effective in causing apoptosis of human osteosarcoma cells ^45,46^. Thus, delphinidin was shown to inhibit metastasis of several cancer types *in vitro* and melanoma (this study) and prostate cancer *in vivo* ^39^, in mouse models. Additional mouse models and human tumor xenografts may be useful to define which cancers are most responsive to delphinidin.

While IP administration of delphinidin was well tolerated and effective in increasing HA levels in multiple tissues, it would be useful to show similar effects with oral administration of delphinidin. It was proposed that encapsulation of delphinidin in small extracellular vesicles could improve bioavailability and enhance its effects ^18^. Another option is Delphinol, an extract from the maqui berry and contains 35% anthocyanins of which 25% is delphinidin. This nutraceutical could be used as an alternative to pure delphinidin for testing in pre-clinical models. Delphinol was used in a clinical trial for patients with glucose intolerance and was shown to lower blood glucose; importantly it had zero adverse effects ^27^. Another clinical trial using Delphinol revealed that its antioxidant properties may reduce oxidized low-density lipoprotein ^45^. In conclusion, pre- clinical mouse models demonstrating inhibition of metastasis by delphinidin together with good safety profile and potentially bioavailable oral administration of delphinidin make it a prime candidate for clinical trials for patients with cancer.

## Material and Methods

### Cell Culture

RM-1 (CRL-3310), E0771 (CRL-3461), B16-F10 Luc2 (CRL-6475-LUC2) were purchased from ATCC. B16-F10 is a sub clone of the murine melanoma cell line from a C57BL/6J mouse. It was generated by injecting mice with B16 tumor cells, collecting and culturing secondary tumor growths, and injecting them into fresh mice, a total of 10 times ^47^. The RM1 cell line was generated he tumor from urogenital sinus Zipras/myc9 retrovirus-injected cells of C57BL/6 embryos ^48^. The EO771 cell line was originally isolated from a spontaneous tumor in C57BL/6 mouse ^49^. Primary C57BL6 skin fibroblasts were isolated as described previously ^50^. All cell lines were cryopreserved within the first five passages and were used within the first 10 passages after thawing. Human fibroblasts are HCA2 originally isolated by Olivia Pereira-Smith, Baylor College of Medicine. All cell lines were cultured at 37 °C, 5% CO_2,_, 3% O_2_ on treated polystyrene culture dishes (Corning). Cancer cell lines were cultured in DMEM, high glucose, GlutaMAX (Gibco Cat # 10566016) supplemented with 10% Fetal Bovine Serum (FBS) (Gibco Cat #10437028) and 1% Penicillin-Streptomycin (Pen Strep) (Gibco Cat #15070063) Gibco. Mouse fibroblasts were culture in Eagle’s Minimum Essential Medium (EMEM) (ATCC Cat# 30-2003) supplemented with 15% FBS and 1% Pen Strep. Cell cultures were periodically tested for mycoplasma contamination with MycoAlert Mycoplasma Detection Kit (Lonza).

### Animals

Wild type C57BL/6 mice were purchased from Charles River. All animal experiments were approved and performed in accordance with ARRIVE guidelines as well as guidelines set up by the University of Rochester Committee on Animal Resources (UCAR), which adheres to FDA and NIH animal care guidelines and reviews all animal protocols prior to approval (UCAR 2017-033). Animals were housed using microisolator technology in SPF conditions in a one-way facility.

### High-throughput screening for Hyaluronidase inhibitor

Fluorescence polarization was utilized to screen hyaluronidase inhibitors. Hyaluronidase (Sigma H4272) and each compound of libraries were incubated in phosphate buffer (300mM, pH 5.35) in black walled black bottom 384 well plates. Fluorescein-labeled HA ^51^ was then added to each well, and the plates were incubated in the dark, humid containers to prevent drying. After 1-hour, fluorescent polarization was measured (TECAN Spark 20M). Compound libraries were obtained from the National Cancer Institute (Approved Oncology Drugs Set, Diversity Set, Natural Products Set and Mechanistic Set).

### HA analysis by gel electrophoresis

HA was purified from conditioned media collected from mouse and human fibroblasts after 24- hour incubation with a dose range of delphinidin. Ten ml of conditioned media was treated with 1 ml Proteinase K (Promega Cat # V3021) in buffer (10 mM Tris-Cl pH8.0, .25 mM EDTA, 100mM NaCl, 0.5% SDS 1mg/ml proteinase K) (Roche Cat #03508838103) at 55 °C for 4 hours to remove proteins. Samples then were mixed by inverting the tubes 60 times after adding 11ml of phenol:chloroform:Isoamyl Alchol (25:24:1) (ThermoFisher Cat# 15593049). Samples were centrifuged with swing rotor at 3500 rpm for 10 minutes. The supernatant was transferred to anew tube and 25 ml of 100% ethyl alcohol (EtOH) was added and mixed by inversion. Samples were centrifuged with swing rotor at 4000 rpm for 1 hour. Supernatant was discarded and 20 ml of 70% EtOH was added and centrifuged for 10 minutes at 4000 rpm. Supernatant was again removed and pellet was allowed to dry for 15 minutes at room temperature. The pellet was dissolved in 2ml PBS and incubated overnight at 4 °C. The following day 20 μl of Triton X- 114 was added to samples and they were vortexed for 5 minutes cooling on ice every 30 seconds. Samples were then incubated at 37°C for 5 minutes before centrifuging at 12,000g for 10 minutes. Supernatant was collected and 5mls of 1M NaCl in 100% EtOH was added and centrifuged at 10,000 g for 15 minutes. Supernatant was then discarded and 5 ml of 70% EtOH was added to pellet and centrifuged at 10,000 g for 15 minutes. Supernatant was then discarded and pellets were dried for 10min at room temperature before dissolving overnight at 4 C in 600ul PBS. 20 ul of each sample was transferred to 2 new tubes and either and treated with 1 U/ml of hyaluronidase from *Streptomyces hyalurolyticus* (Sigma-Aldrich Cat # H1136) or no treatment and then incubated overnight at 37°C. This protocol was previously published ^52^. For gel electrophoresis, a 0.8% Seakem Gold Agarose (Lonza Cat#50150) gel/TBE was kept at 55°C for 15 minutes before pouring into gel tray. After 2 hours, the gel was pre-run in TBE buffer at 35V for 2 hours. 10ul of each sample were then diluted in 6X loading buffer. Ladders and samples were then loaded and run at 23V for 30 minutes and then 35V for 3 hours or until yellow dye runs out of gel. Gel was then stained overnight in the dark at room temperature in 0.005% Stains-All (Sigma Cat# E9379) prepared in 50% EtOH. The following day the gel was washed twice in 10% EtOH for 10 minutes and then washed a third time for 4 hours or overnight. The gel was then imaged using digital photography. The gel was scanned using Bio Rad Gel Imager and quantitative analysis of HA was assessed using GelQuant.Net (http://biochemlabsolutions.com/GelQuantNET.html). The original gel image is no longer available as the bio rad gel imager had to be replaced and the stored images were lost. Only the top of the gel showing the wells was cropped out of the image used in figure 1. Full gel images from supplemental figure 1 are included in the supplemental data.

### *In vivo* delphinidin treatment

Three to six-month-old female C57Bl/6 (n=10) mice were treated with either 50 mg/kg (n=4) or 100 mg/kg (n=3) of delphinidin chloride (Cayman Cat# 11012). The delphinidin was dissolved in 10% Dimethyl sulfoxide (DMSO) and 90% sterile PBS that was used as a vehicle control (n=3). Intraperitoneal (IP) injections of either delphinidin or vehicle control (200 were administered three times a week (n=3 Vehicle, n=4 50 mg/kg, and n=3 100 mg/kg) for three weeks before the mice were sacrificed by CO2 euthanasia followed by secondary cervical dislocation. The following tissues, lung, kidney, spleen, heart, skin, muscle, duodenum, jejunum and colon were collected fixed in 10% formalin for 48 hours before being embedded in paraffin.

### Ear transplantation of B16F10 *in vivo*

B16-F10 Luciferase expressing cells were suspended in PBS at 10×10^6^/ml and 50 l of suspension were injected between the skin and cartilage on the dorsal side of the ear for a total of 5X10^5^ cells injected per mouse. C57/BL6 mice at approximately 8 months of age were used. Male (n=20) and female (n=20) mice (C57/BL6, n=40) were anesthetized by isoflurane during transplantation. The tumor take was 60% and 25% for the delphinidin and vehicle groups respectively. One week prior to tumor cell inoculation one cohort of mice were treated with delphinidin 50 mg/kg diluted in 10% DMSO and 90% PBS by IP injection (200 µl) three times a week. Treatment was continued 3 times a week for the duration of the experiment the other cohort of mice were injected with the vehicle for the delphinidin (200ul 90% PBS 10% DMSO). Outcome was defined as time to detectable metastasis. Endpoint was when primary tumor sizes reached (2 mm^2^) at which point mice were sacrificed by CO2 euthanasia followed by secondary cervical dislocation. Due to the study design for treating and imaging the mice randomization and blinding was not possible and no confounders were controlled for. Bioluminescence was measured once weekly as a correlate of tumor growth and detection of metastases (IVIS, Perkin Elmer, Waltham, Massachusetts).

### HABP staining

Slides with 2 sequential 5mm paraffin sections (n=3 for each animal) washed two times with xylenes 10 minutes each then rehydrated in ethanol 2 min each (100%, 100%, 90%, 70% 50%). Rinsed in 1X PBS 3 times, 5 min each. Tissue sections circled with PAP pen and incubated for an hour at room temperature in (100-200 ml) blocking solution (10% normal goat serum (NGS) (Fisher Cat# 50062Z) in 1X PBS). The blocking solution was discarded, and slides were incubated in biotinylated hyaluronic acid binding protein (HABP) (Amsbio Cat# AMS.HKD-BC41) at a 1:200 dilution in 10% NGS overnight at 4 C with gentle rocking in a humidified chamber. Negative controls consisted of two sections from each group that were incubated for 2 hours at 37 C with 1 mg/ml hyaluronidase (Sigma Cat# H3884) in 20 mM sodium phosphate buffer, with 77 mM sodium chloride, 0.01% BSA, pH 7.0 before HABP treatment. The following day slides were washed three times in 1X PBST (0.01% Tween 20) 10 min each wash followed by 2 rinses in 1X PBS. Sections were incubated with streptavidin alexa fluor 488 (Invitrogen Cat # S32354) 1:1000 dilution in 10% NGS in the dark at room temperature for 1 hour. Slides were washed 3X in PBST 10 minutes each and rinse 2X in 1X PBS 1 min each. Cover slips were mounted with Vectashield plus antifade mounting medium with DAPI (Vector laboratories Cat#H2000). Slides were imaged with fluorescence microscopy (20X), three representative images were analyzed from each tissue section (2 tissue sections from each sample) using Imaris software version 9.8 Oxford Instruments (https://imaris.oxinst.com/) to determine maximum fluorescent intensity. Background fluorescence from negative control, hyaluronidase treatment, was subtracted from all samples.

### Proliferation Assays

Cells were seeded in triplicate at 2X10^4^ -5X10^4^ depending on the cell type in white walled white bottom 96 well plates and incubated overnight. On the following day cells were treated with 5 mM delphinidin (stock 30 mg/ml) diluted in media before adding to cells. Hyaluronidase (Sigma Cat # H1136) was added with delphinidin in media before adding to cells. Plates were analyzed at 24 and 48 hours post delphinidin treatment with Cell Titer-Glo 2.0 cell viability assay. (Promega Cat#G9241). 100 ml of cell titer-glo was added to each well and incubated shaking in the dark at room temperature for 10 minutes before reading luminescence using a plate reader.

### Wound Healing Assay

Three well silicone inserts (Ibidi Cat#80369) were placed with sterile forceps into individual wells of a 24 well plate and seeded with 70 ml of cell suspension 5X10^5^ cells/ml. After 24-48 hours when the cells reached confluency the inserts are removed with sterile forceps and the wells were filled with 1 ml of cell culture media either with or without delphinidin. Cells are then immediately imaged and then imaged again at 8-20 hours depending on the cell type. Using image j the area of the gap is measured at time 0 and at 8-20 hours.

### Migration and Invasion Assays

Cells were seeded and allowed to grow to 80% confluence then serum starved overnight. On the next day the cells were seeded (2 × 10^5^ cells) into upper chambers of 8 mM 24-well transwell membrane assay system (Corning Cat#3428). Lower chambers were prepared either 650 µl serum free (-control), 1% serum (+control) or 1% serum with delphinidin or 1% serum with delphinidin and hyaluronidase. All conditions were performed in triplicate and incubated for 5 h at 37 °C for migration assays. For invasion assays the upper chamber membranes were coated with Matrigel Basement Membrane Matrix (Corning Cat#354234). The Matrigel was thawed on ice, diluted with serum free media to a concentration of 200 mg/ml, mixed by pipetting and 100 ml added to each upper well and then incubated for at least 1 hour at 37 C to let the gel form. The cells were then seeded in the Matrigel treated wells. After incubation, upper chambers were rinsed in DI water followed by 1× PBS. Upper chambers then fixed in pure ice-cold methanol at −20 °C for 20 min. The chambers were rinsed in DI water followed by 1× PBS, and non-migrated cells removed from the inner membrane by gentle scrubbing 3X with cotton tip applicators. After scrubbing membranes are rinsed again in DI water before being stained with hematoxylin for 5 minutes and incubated in 1X PBS to blue then cells. Upper chambers are then rinsed 3X in DI H20 before dehydrating in 100% ethanol for 3 minutes. The chambers were allowed to air dry for 10 minutes before using a scalpel to cut out the membrane. Membranes were fixed to microscope slides bottom up with permount (Fisher Cat# sp15100). Three fields of view from each membrane were acquired using brightfield microscopy (10X) and the number of migrated cells per field counted using image j.

### Statistical Analysis

T-tests or analysis of variance (ANOVA) and post hoc Tukey’s multiple comparison tests were used to calculate statistical significance (≤0.05) among groups. Samples sizes of n≥3 was used for all statistical calculations. Error bars are shown as standard error from the mean (SEM). These statistical analyses were calculated with the Graph Pad Prisim 10.0, GraphPad Software, Boston, Massachusetts USA, (www.graphpad.com).

## Data Availability

All the data generated or analyzed during this study is available from the corresponding author on reasonable request.

## Acknowledgements

This work was supported by grants from US National Institutes on Aging to VG and AS and a grant from VitaDao to V.G.

## Contributions

V.G., A.S, T.T. and J.M were responsible for the study conception as well as the design and development of methodology. J.M, T.T and V.P. acquired data. V.G, A.S, T.T, G.T. and J.M were responsible for analysis and interpretation of data. V.G., A.S., G.T., N.G., M.J., T.T and J.M were responsible for manuscript writing and editing.

