## Supplemental Figures for "Hyaluronidase inhibitor delphinidin inhibits cancer metastasis"

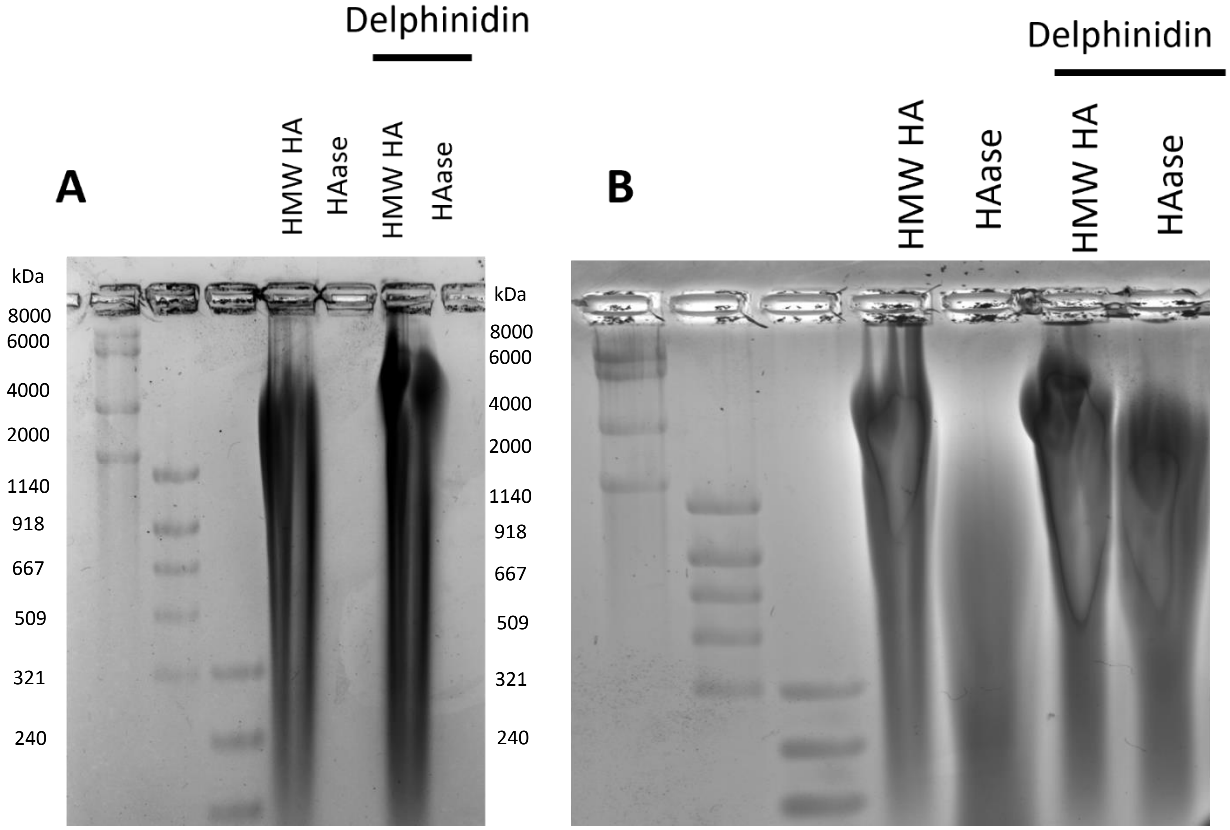


**Supplemental Figure 1:** Delphinidin increases HMW HA levels in cultured cells and inhibits degradation by HAase in cell culture media***: A.*** Delphinidin inhibits HAase from degrading HMW HA in cell culture media. Exogenous HA was extracted from cell culture media of B16-F10 cancer cells and treated with HAase overnight as a negative control. **B.** A diluted stock of HAase was used to degrade HA. The sample treated with delphinidin was not degraded as much as the sample not treated with delphinidin.

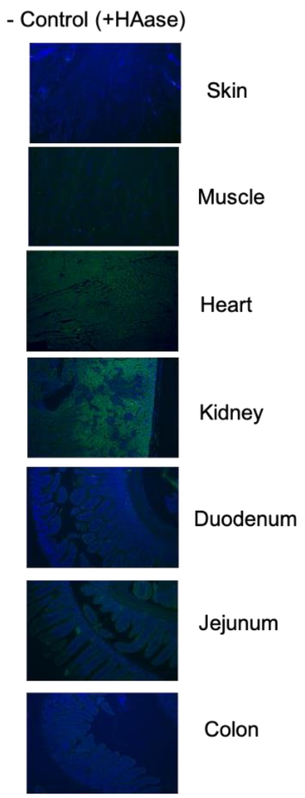


**Supplemental Figure 2: HAase treated mouse tissues before HABP staining (- Control) :** As a negative control mouse tissues were treated with 1 mg/ml hyaluronidase for 2 hours before incubating in HABP for staining. Mean background fluoresce from the negative control tissues was subtracted from all samples when calculating mean fluoresce intensity for Figure 2.

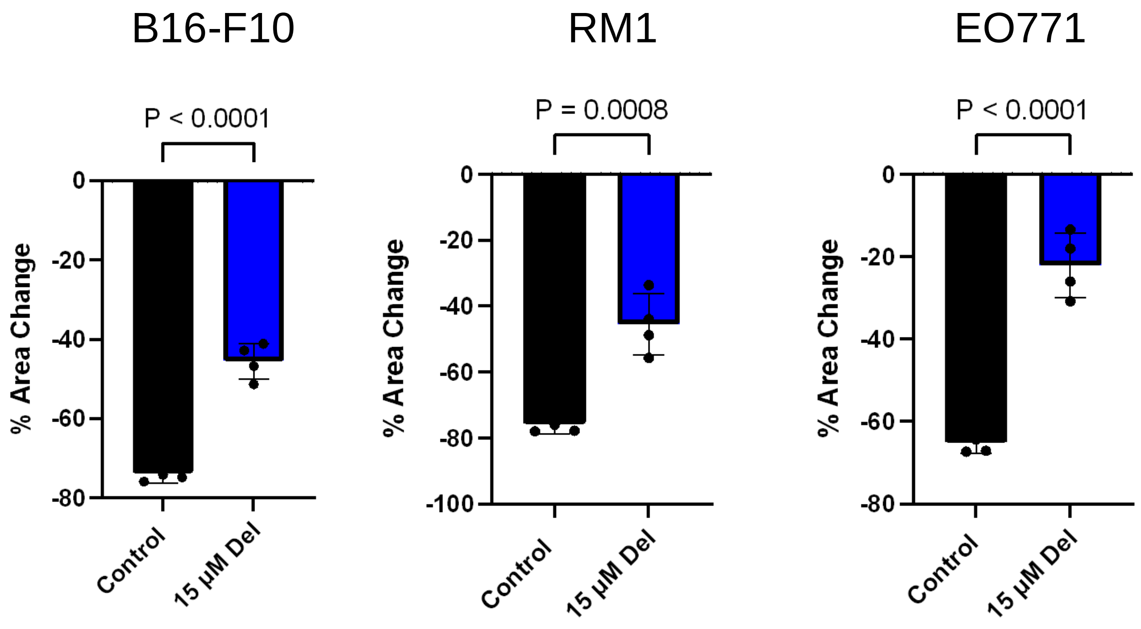


**Supplemental Figure 3:** Delphinidin inhibits migration of cancer cells. 15 µM Delphinidin inhibits wound migration (rate of gap closure) when compared to non-treated control of all three cancer cell lines in an 8-15-hour migration assay. A. B16-F10 B. RM1 C. EO771. Data was collected as technical triplicates and statistical analyses were generated by unpaired t-test error bars are represented as mean with SEM.

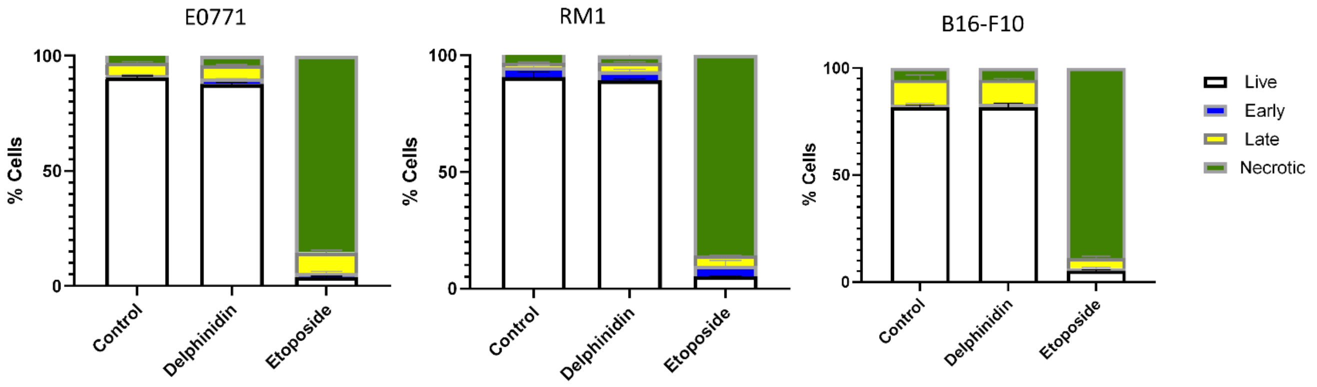


**Supplemental Figure 4:** Annexin 5 staining of cancer cells: EO771, RM1 and B16-F10 were treated with delphinidin (30ul) or etoposide (50ul) for 24 hours before staining. There was very little apoptosis detected in any of the cell lines. Data was collected as technical triplicates.

**Supplemental Information:**

**Uncropped Gel Images (Supplemental Figure 1):**

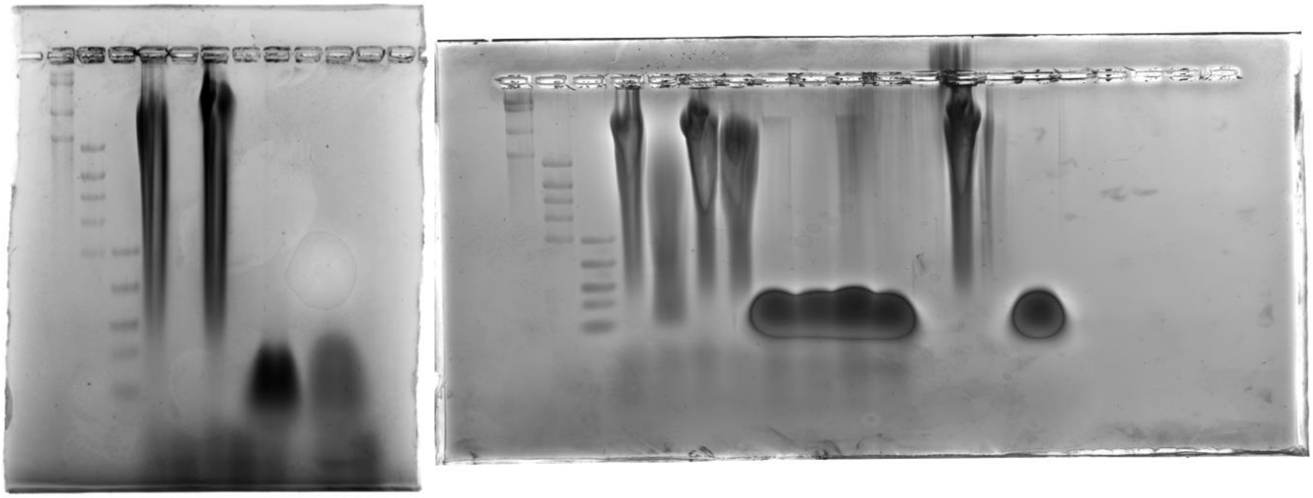
